# ULK1 and ULK2 are less redundant than previously thought: Computational analysis uncovers distinct regulation and functions of these autophagy induction proteins

**DOI:** 10.1101/2020.02.27.967901

**Authors:** Amanda Demeter, Mari Carmen Romero-Mulero, Luca Csabai, Márton Ölbei, Padhmanand Sudhakar, Wilfried Haerty, Tamás Korcsmáros

**Affiliations:** Earlham Institute, Norwich Research Park, Norwich, NR4 7UZ, UK; Quadram Institute Bioscience, Norwich Research Park, Norwich, NR4 7UQ, UK; Faculty of Biology, University of Seville, Seville, 41012, Spain; Eötvös Loránd University, Budapest, 1117, Hungary; Department of Chronic Diseases, Metabolism and Ageing, KU Leuven, Leuven, Belgium

## Abstract

Macroautophagy, the degradation of cytoplasmic content by lysosomal fusion, is an evolutionary conserved process promoting homeostasis and intracellular defence. Macroautophagy is initiated primarily by a complex containing ULK1 or ULK2 (two paralogs of the yeast Atg1 protein). Deletion of ULK1 is sufficient to interrupt autophagy, while ULK2 seems expendable. To understand the differences between ULK1 and ULK2, we compared the human ULK1 and ULK2 proteins and their regulation. Despite the high similarity in their enzymatic domain, we found that ULK1 and ULK2 have major differences in their post-translational and transcriptional regulators. We identified 18 ULK1-specific and 7 ULK2-specific protein motifs serving as different interaction interfaces. We identified three *ULK1*-specific and one *ULK2*-specific transcription factor binding sites, and eight sites shared by the regulatory region of both genes. Importantly, we found that both their post-translational and transcriptional regulators are involved in distinct biological processes - suggesting separate functions for ULK1 and ULK2. For example, we found a condition-specific, opposite effect on apoptosis regulation for the two ULK proteins. Given the importance of autophagy in diseases such as inflammatory bowel disease and cancer, unravelling differences between ULK1 and ULK2 could lead to a better understanding of how autophagy is dysregulated in diseased conditions.

## Introduction

Macroautophagy, hereafter referred to as autophagy, is a lysosome-dependent intracellular metabolic process. Autophagy is highly conserved in all eukaryotic cells, and contributes to maintaining energy homeostasis, generating nutrients following starvation and is crucial in promoting cell survival. Autophagy is present in cells at a basal level but it is also activated at a higher level as a response to stress^1,2^.

Upon autophagy induction, an isolation membrane (phagophore) sequesters a small portion of the cytoplasm, including soluble materials and organelles, to form the autophagosome. The autophagosome then fuses with the enzyme-containing lysosome to become an autolysosome and degrades the materials contained within it^1^. As observed first in yeast, induction of autophagy is governed by the induction complex formed by Atg proteins: Atg1-Atg13-Atg17-Atg31-Atg29. In mammalian cells, the complex is composed of Atg1 homologs (ULK1 or ULK2), the mammalian homolog of Atg13 (ATG13), RB1-inducible coiled-coil 1 (RB1CC1/FIP200) and ATG101^3,4,5^. Induction of autophagy is tightly regulated through the protein mechanistic target of rapamycin complex 1 (mTORC1): whilst complex-associated, mTORC1 phosphorylates ULK1/2 and ATG13, resulting in their inactivation. Nevertheless, when cells are treated with rapamycin or starved of nutrients, mTORC1 becomes separated from the induction complex, resulting in dephosphorylation of these proteins and consequent autophagy induction^6^. Interestingly, this type of autophagy induction is specific to higher eukaryotes and some of the protist, however, as we previously demonstrated, non-unikont protists (such as *Toxoplasma spp.*, *Leishmania spp.* and *Plasmodium spp.*) lack the Atg1 complex, and induce their autophagy in different ways^7^.

The most studied component of the induction complex is the yeast *Atg1* and its homologs. There are five *Atg1* orthologs in the human genome: *ULK1, ULK2, ULK3, ULK4,* and *STK36*, all codes proteins containing a kinase domain (Figure 1)^2,8^. Out of these genes, the protein product of *ULK1* and *ULK2* shows high similarity along the whole length of the protein (98% query cover but only 52% identity)^10^ and in their kinase domain (100% query cover with 78.71% identity)^2^. *ULK1* and *ULK2* genes code for serine/threonine protein kinases that consist of a conserved N-terminal kinase domain (catalytic domain), a central serine-proline rich region (PS), and a C-terminal interacting domain (CTD)^9^. ULK1 and ULK2 are best known to be involved in the induction of autophagy^10,11^, and are often mentioned interchangeably, however, there is growing evidence for functional differences. For instance, Seung-Hyun *et al.* found that while both proteins have shared functions in autophagy, they have opposing roles in lipid metabolism^10^. ULK1 and ULK2 were also shown to be part of different molecular complexes, possibly resulting in having independent functions or mode of regulation. It was specifically found that following amino acid starvation, in contrast to ULK1, binding of ULK2 to membranes was increased^12^. In most cell lines studies, loss of *ULK1* is sufficient to interrupt autophagy, and *ULK2* was thought to have a redundant function. On the other hand, in mice studies, the knockout of both *ULK1* and *ULK2* was needed to show the neonatal lethality, which was seen also upon the loss of other core autophagy genes such as *ATG5* and *ATG7*^*13–15*^. During erythroid differentiation, *ULK1*, but not *ULK2* was found to be upregulated, and deletion of *ULK1*, hence autophagy deficiency resulted in delayed clearance of ribosomes and mitochondria in reticulocytes^13^. Lee and Tournier also showed that the function of ULK2 is more likely to compensate for the loss of ULK1 but in a cell-type specific manner^2^. Some speculations indicate a dominance of either of the two ULK proteins determined by differential tissue expression levels^15,16^.

**Figure 1:**
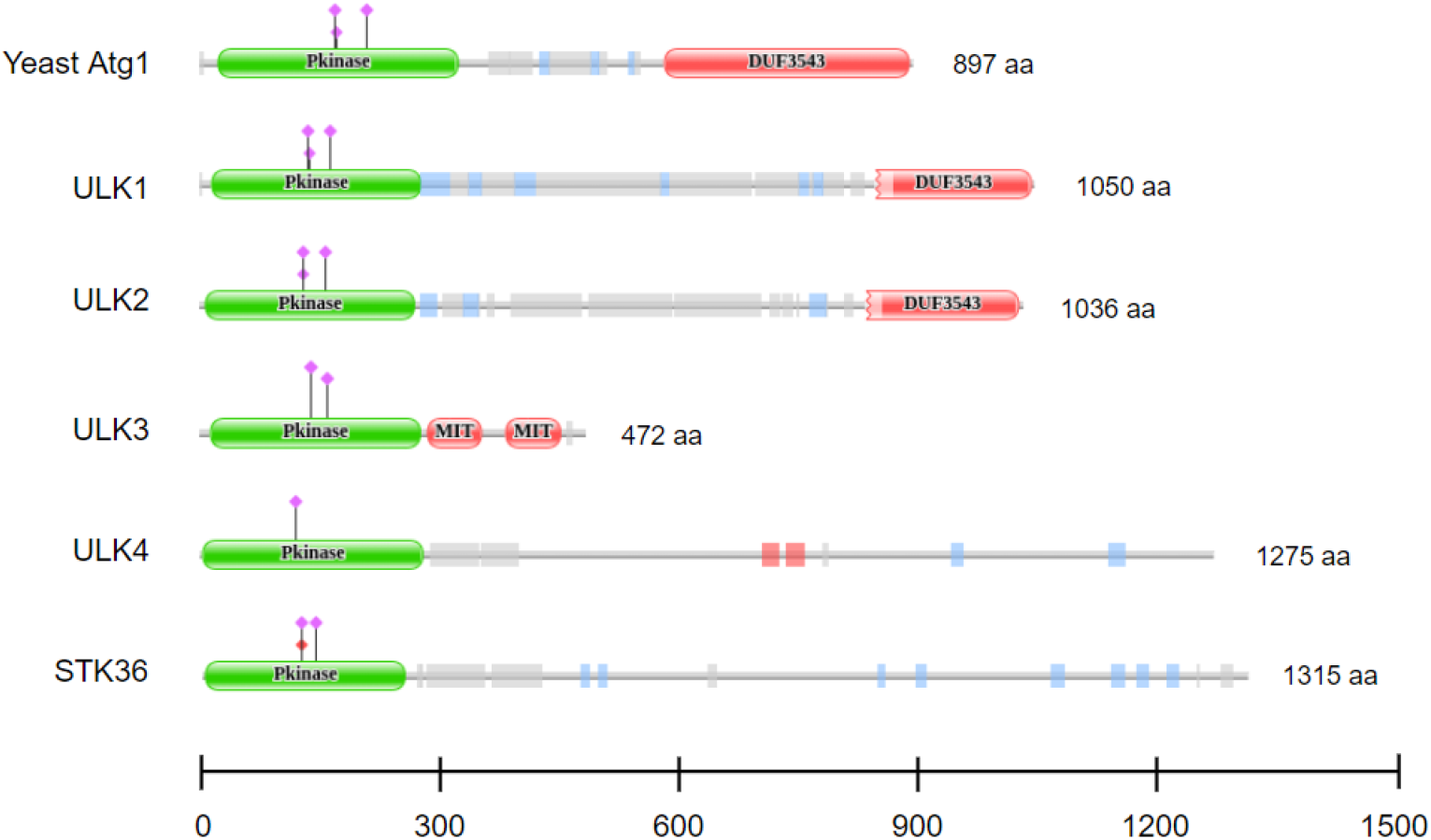
Domain structure of the yeast Atg1 protein and its human homologs. All Atg1 homologs have an N-terminal kinase domain, while only Atg1, ULK1 and ULK2 share the C-terminal DUF3543 domain. The domain structure of the proteins were downloaded from the.Pfam website (http://pfam.xfam.org/^30^).

Prompted by the evidence for the different roles of ULK1 and ULK2, we aimed to compare the two human paralogs on a systems-level. We show that ULK1 and ULK2 are indeed different in their protein binding motifs and in their promoter sequences, harbouring binding sites for different transcription factors. They also have different experimentally validated post-translational and transcriptional regulators and their respective interactors are important in distinct biological processes. The balance between these two homologs and their separate processes can therefore be crucial in diseases, especially when ULK1, ULK2 or their interactors are differentially expressed.

## Results

### The duplication of the yeast *Atg1* happened at the base of the Chordates resulting in the most similar homologs, *ULK1* and *ULK2*

The multiple sequence alignment of the yeast Atg1 protein and its human orthologs shows that the human ULK1 and ULK2 are the most similar to each other (Figure 2). We obtained similar results with alignment of the kinase domain and alignment of the complete amino-acid sequences.

**Figure 2:**
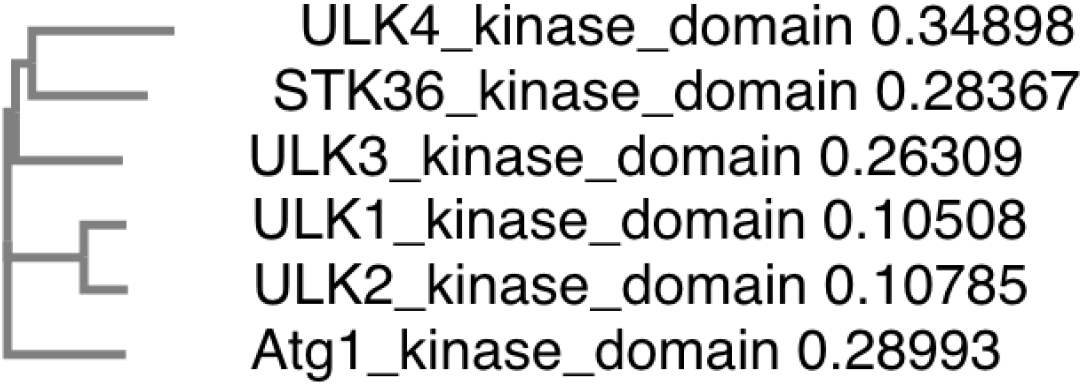
Phylogenetic tree of kinase domains of the yeast Atg1 protein and its human homologs. Based on multiple sequence alignment of the kinase domains ULK1 and ULK2 are the most similar to each other (similar results were obtained with multiple sequence alignment of the complete amino-acid sequences, see Supplementary figure S1). The protein sequences were downloaded from Uniprot (https://www.uniprot.org/)^32^ and the protein alignments were carried out with MUSCLE (https://www.ebi.ac.uk/Tools/msa/muscle/)^33^

We searched for the orthologs of ULK1 in other species to determine the first duplication event that gave rise to *ULK1* and *ULK2* (in humans). On Figure 3, we visualize the duplication event which happened at the base of the Chordates, after the split of Urochordates (Tunicates, around 500 million years ago) from the Euteleostomes or “bony vertebrates”. As expected, in Deuterostomes and in *Ciona intestinalis* we can find only one ortholog of the human *ULK1* gene, but in the rest of the Chordates most taxa have two paralogs.

**Figure 3:**
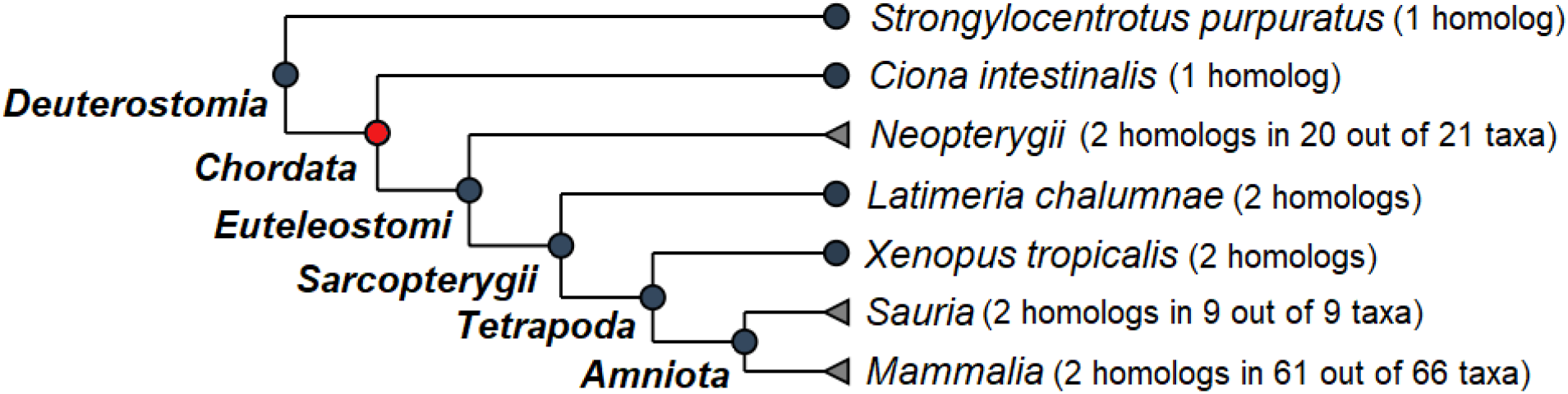
Phylogeny of the human *ULK1 gene*. Duplication of the *ULK1* gene happened at the base of the Chordates, after the split of *Ciona intestinalis* from the Euteleostomes or “bony vertebrates”. The dendrogram was adapted from the OMA orthology browser (https://omabrowser.org/)^31^.

### ULK1 and ULK2 have specific protein-protein interaction partners

We collected the experimentally verified protein interactors of ULK1 and ULK2 (termed as first neighbours) from databases and from the literature by manual curation. Out of the 25 ULK1 first neighbours and the 35 ULK2 first neighbours, 11 and 21 are specific to ULK1 and ULK2, respectively.

On Figure 4, we visualised the experimentally validated directed interactions between the ULK proteins and their first neighbours. We grouped the neighbours based on being upstream or downstream from their respective ULK protein and based on being involved in single or bidirectional interaction. We also show that most of the first neighbours are predicted to bind to both ULK1 and ULK2, indicating a potential study bias in the literature data. The presence of the common downstream interactors in the prediction is possibly due to the domain similarity of the ULK homologs.

**Figure 4:**
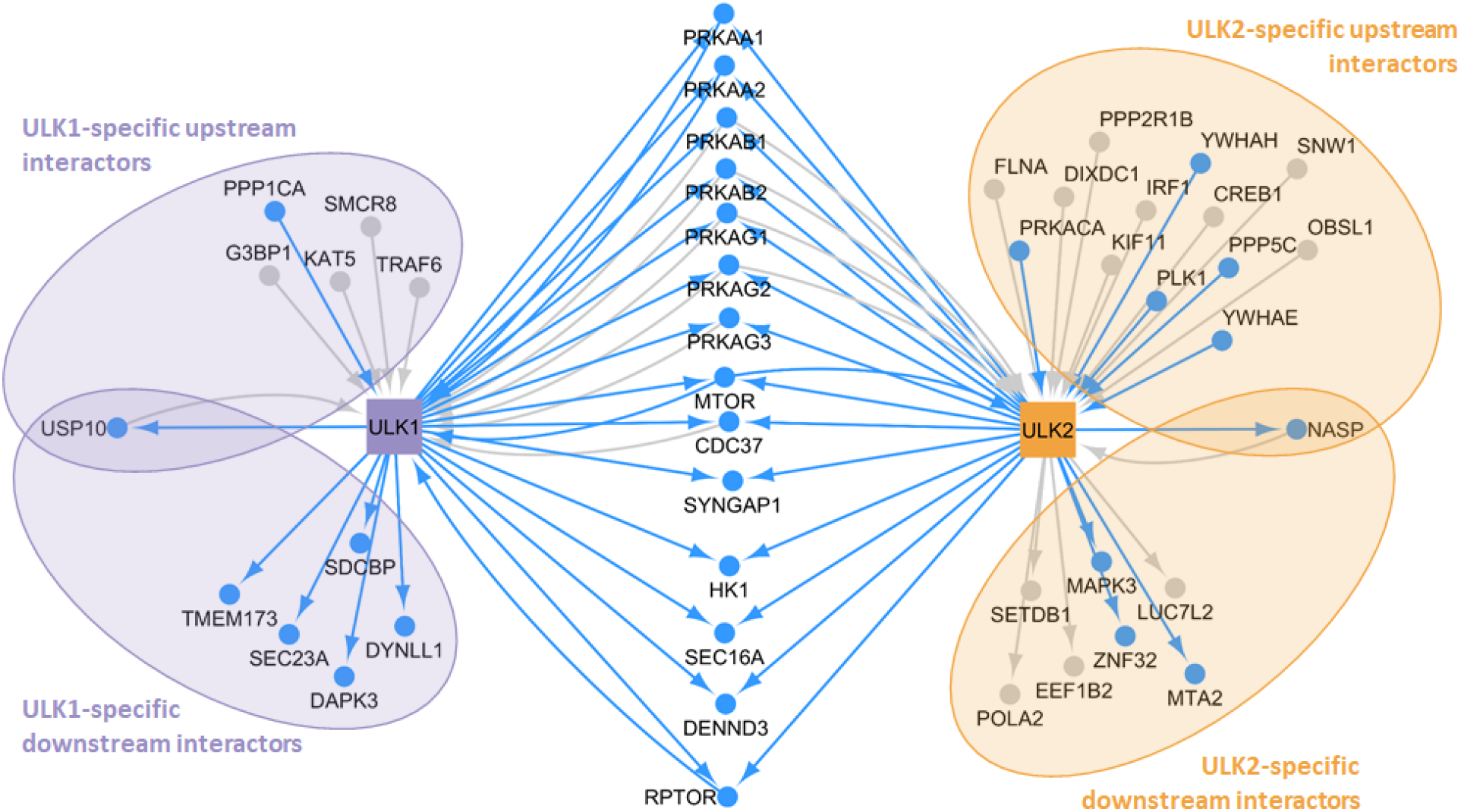
Experimentally validated protein-protein interactions of ULK1 and ULK2 collected from databases and from the literature by manual curation. Blue nodes and edges represent those experimentally identified interaction partners and interactions, respectively, that we found to be supported by a protein binding motif analysis (done with the Pfam^30^ and ELM resource^34^). Grey nodes and edges show experimentally identified ULK-specific interaction partners and interactions, respectively for which the protein motif analysis did not provide potential binding interfaces.

As both ULK1 and ULK2 have a kinase domain with 78.71% identity (Figure 1), we hypothesized that the explanation for having specific interactors could be primarily due to exhibiting different protein motifs. Based on structural information, we found that beside sharing 37 motifs, ULK1 and ULK2 have 18 and 7 unique motifs, respectively (Table 1 and Table 2). Amongst the ULK1-specific motifs, we identified sites for caspase cleavage, sister chromatid separation and binding to inhibitor of apoptosis proteins, which is in favour of promoting apoptosis. On the other hand, amongst the ULK2-specific motifs there are sites for deubiquitination and interaction with TNF cytokine receptors.

**Table 1:** ULK1-specific motif details from the ELM resource (http://elm.eu.org/)^34^. The list and annotated functions of 18 ULK1-specific motifs.

| ULK1-specific motif name | Functional site class | Function |
| --- | --- | --- |
| CLV_C14_Caspase3-7 | Caspase cleavage motif | Caspase-3 and Caspase-7 cleavage in apoptosis. |
| CLV_Separin_Metazoa | Separase cleavage motif | Sister chromatid separation in the metaphase-to-anaphase transition in cell division |
| DEG_APCC_DBOX_1 | APCC-binding Destruction motifs | Proteasome-dependent degradation of proteins in a cell cycle dependent manner by ubiquitination mediated by APC/C. |
| DEG_COP1_1 | COP1 E3 ligase binding degron motif | Proteasome-dependent degradation of proteins by ubiquitination. |
| DOC_ANK_TNKS_1 | Tankyrase-binding motif | PARsylation of target proteins by Tankyrase regulating their ubiquitylation, stability and function. |
| DOC_CyclinA_RxL_1 | Cyclin docking motif | Protein phosphorylation by cyclin/Cdk complexes involved in biological processes eg. stress response, cell division checkpoints |
| DOC_MAPK_FxFP_2 | MAPK docking motifs | Increased binding affinity to specific interactors of the MAPK cascade, regulators of cellular signaling. |
| LIG_Actin_WH2_2 | Actin-binding motifs | Regulation of the actin filaments assembly influencing cellular functions such as the control of cell shape or cellular transport. |
| LIG_BIR_III_2 | IAP-binding motif (IBM) | Binding to Inhibitor of Apoptosis Proteins (IAPs) leading to apoptosis promotion. |
| LIG_CtBP_PxDLS_1 | CtBP ligand motif | Interaction and recruitment into nuclear complexes of C-terminal binding proteins, repressors of transcription. |
| LIG_EVH1_1 | EVH1 ligands | Binding to signal transduction class I EVH1 domains, influencing signal transduction, such as re-organization of the actin cytoskeleton. |
| LIG_GBD_Chelix_1 | GTPase-binding domain (GBD) ligand | Binding to the GTPase-binding domain (GBD) in WASP and N-WASP, thus preventing Arp2/3-dependent activation of actin polymerization. |
| LIG_PDZ_Class_2 | PDZ domain ligands | Recognition of short sequences at the carboxy terminus of target proteins, developing a variety of biological processes including cell signalling and synapse. |
| LIG_Pex14_2 | Pex14 ligand motif | Binding to Pex14, a key protein in peroxisomal import. |
| LIG_SH3_1 | SH3 ligand | Interactions involved in biological processes: signal transduction, cytoskeleton organization, membrane traffic or organelle assembly. |
| LIG_TYR_ITIM | Immunoreceptor tyrosine-based motif | Recruitment and activation of a protein tyrosine phosphatase that regulates intracellular response of T, B and natural killer cells. |
| LIG_WW_3 | WW domain ligands | Recognition of proteins that contain the motif PPR, being involved in many cellular processes such as ubiquitin-mediated protein degradation and mitotic regulation and several human diseases. |
| MOD_NEK2_2 | NEK2 phosphorylation site | NEK2 phosphorylation motif which can involve many cell cycle-related functions, including cell cycle progression or chromatin condensation. |

**Table 2:** ULK2-specific motif details from the ELM resource (http://elm.eu.org/)^34^. The list and annotated functions of 7 ULK2-specific motifs.

| ULK2-specific motif name | Functional site class | Function |
| --- | --- | --- |
| DOC_MAPK_DCC_7 | MAPK docking motifs | Interaction towards the ERK1/2 and p38 subfamilies of MAP kinases, regulators of cellular signaling. |
| LIG_APCC_ABBA_1 | APCC activator-binding ABBA motif | Regulation of APC/C activity in ubiquitination for proteasome-dependent degradation in a cycle-dependent manner. |
| LIG_CaM_IQ_9 | Helical calmodulin binding motifs | Primary receptor of intracellular Ca<sup>2+</sup> capable of responding to a wide range of calcium concentration and translates the Ca<sup>2+</sup>-signal into cellular processes |
| LIG_EF_ALG2_ABM_2 | ALG-2 EF hand ligand | Binding to ALG-2 protein in a Ca<sup>2+</sup>-dependent manner. ALG-2 has been implicated in ER-stress-induced apoptosis, cell cycle progression, the endosomal pathway, and cancer. |
| LIG_SH2_GRB2like | Phosphotyrosine ligands bound by SH2 domains | Binding to specific motifs containing a phosphorylated tyrosine residue, propagating the signal downstream promoting protein-protein interaction and/or modifying enzymatic activities. |
| LIG_TRAF6 | TRAF6 binding site | Initiation of intracellular signaling caused by members of the tumor necrosis factor receptor (TNFR) superfamily, and direct interaction with intracellular regions of various TNF cytokine receptors. |
| LIG_WW_1 | WW domain ligands | Recognition of proteins that contain the motif PPXY, being involved in many cellular processes such as ubiquitin-mediated protein degradation and mitotic regulation and several human diseases. |

### The first neighbour interactors of ULK1 and ULK2 are involved in different biological processes

We analysed the shared biological processes within all experimentally verified protein interactors of ULK1 and ULK2, respectively. Based on a Gene Ontology Term Finder analysis^17^, we found that beside the common biological functions, the interactors also seem to be responsible for specific processes. Based on our analysis, interactors of ULK1 share more functions that are related to catabolism (Bonferroni corrected *P*-value for the hypergeometric distribution <0.0001) with KAT5, USP10, SDCBP being involved specifically in ubiquitin-dependent degradation. Meanwhile, ULK2 has interactors relevant in general metabolic pathways like organic substance metabolic process (corrected *P*-value <0.0001) and regulation of macromolecule metabolic process (corrected *P*-value <0.0001) (Figure 5).

**Figure 5:**
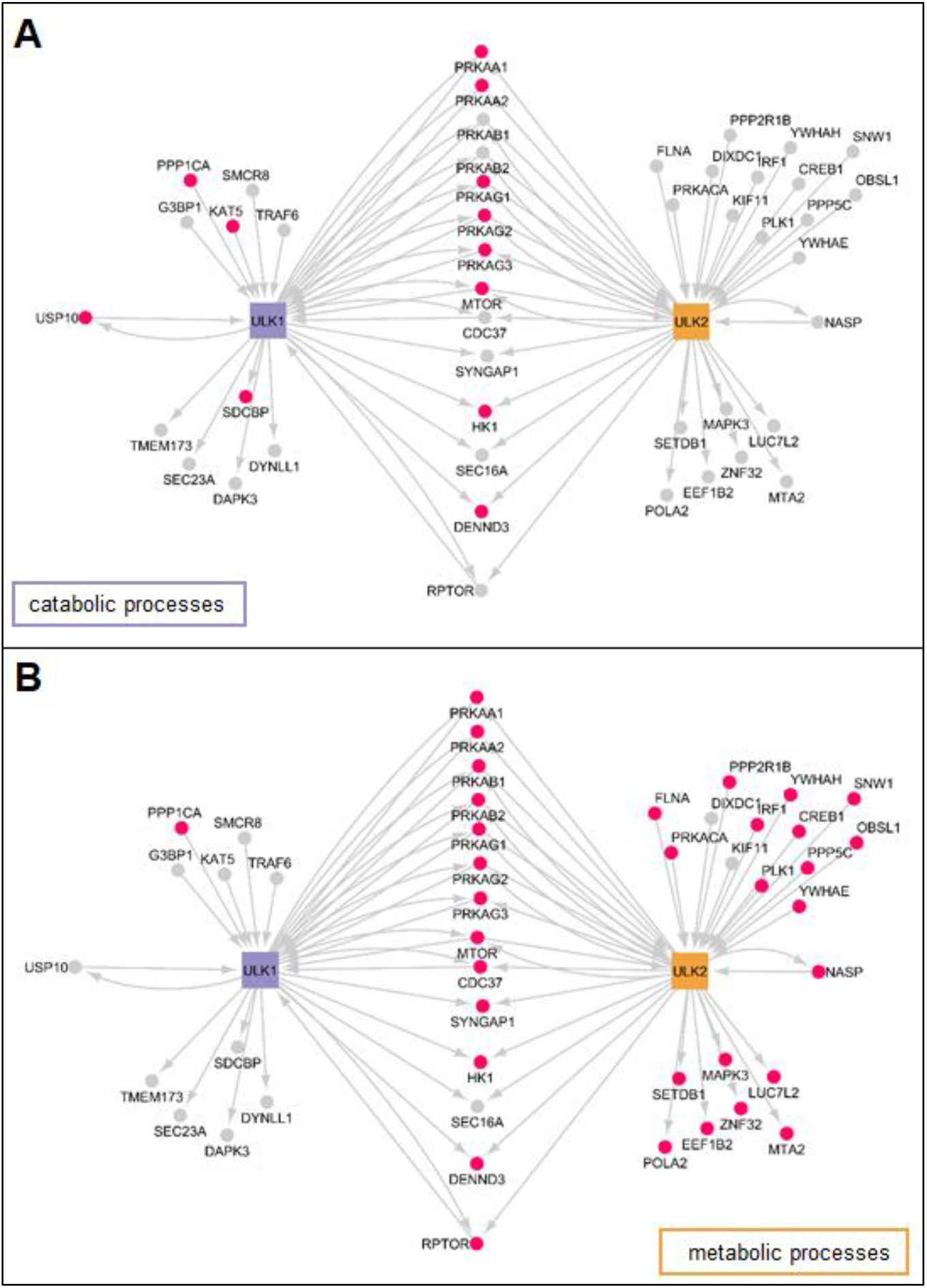
The protein-protein interaction partners of ULK1 and ULK2 and two biological processes predominantly being associated to ULK-specific protein interactors. The biological processes are labelled on each network, interactors involved in the respective biological process are highlighted with pink nodes. A) catabolic processes; B) metabolic processes.

### *ULK1* and *ULK2* have specific transcriptional regulators, which are also involved in different biological processes

We searched for experimentally verified transcription factors of *ULK1* and *ULK2* in databases and the literature by manual curation. Out of the 14 regulators of *ULK1* and 13 regulators of ULK2, six and five are specific, respectively. These transcription factors are likely to regulate the two *ULK* homologs differently. To extend the analysis, we predicted potential transcription factor binding sites (TFBSs) in the promoter regions of both *ULK* genes. We found that six transcription factors (TP53, ATF4, ESR1, HIF1A, CEBPA, FOXP1) that were experimentally identified as *ULK1 or ULK2*-specific, have actually binding sites along the other *ULK* gene as well, indicating potential new regulatory connections for experimental validation (Figure 6). Those transcription factors whose TFBS was not found in the promoter region of *ULK1* or *ULK2* but previous experimental data showed the regulatory connection, are probably regulating the ULK gene’s expressions through more distant regulatory regions (e.g. in enhancer regions).

**Figure 6:**
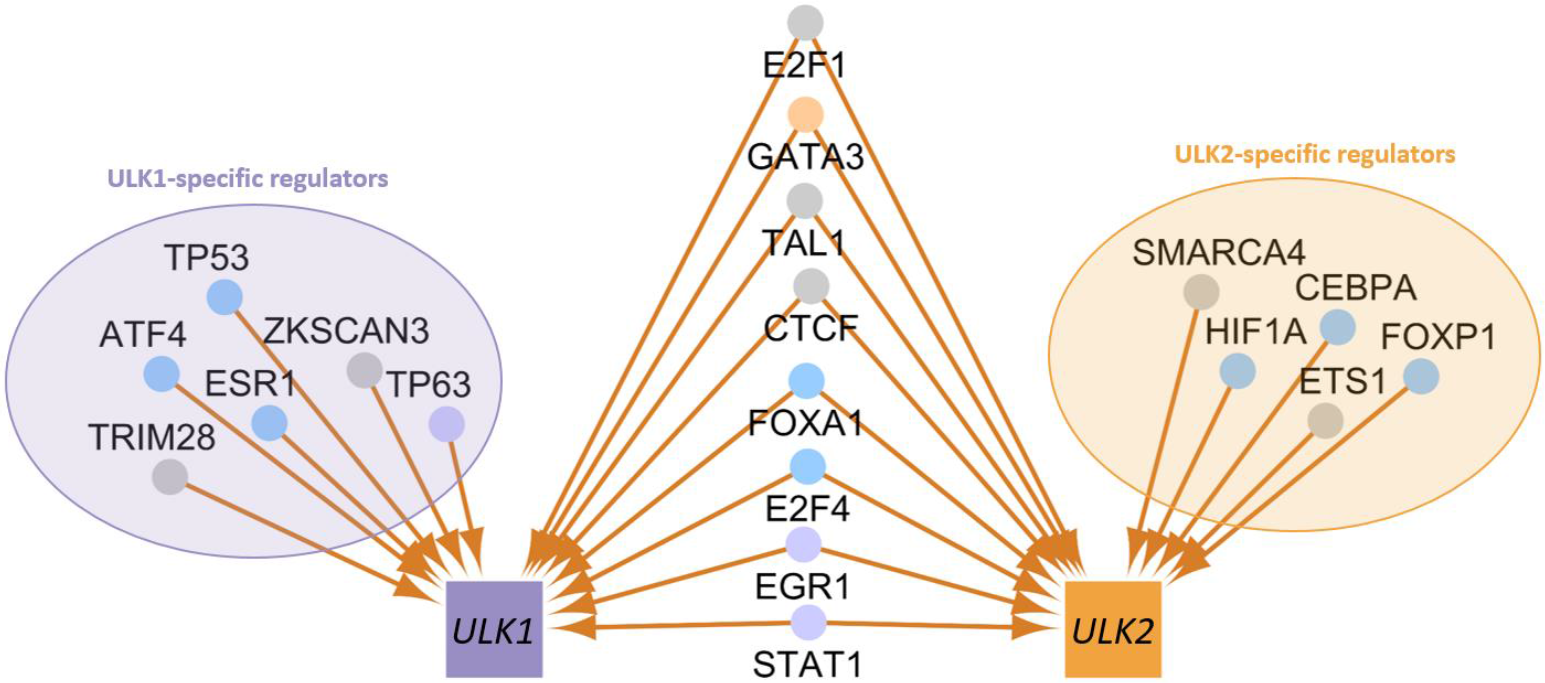
Experimentally validated transcriptional regulators of *ULK1* and *ULK2*. The nodes of the transcription factors are coloured based on the result of the transcription factor binding site (TFBS) analysis: Light purple: predicted TFBS supporting the experimental data was found on the *ULK1* promoter sequence; blue: predicted TFBS was found on the *ULK1* and *ULK2* promoter sequences; light orange: predicted TFBS was found on the *ULK2* promoter sequence; grey: no predicted TFBS was found.

We also analysed the biological processes in which the transcription factors of *ULK1* and *ULK2* are involved. Based on a Gene Ontology Term Finder analysis^17^, we found that transcription factors of ULK1 are significant in stress response (Bonferroni corrected *P*-value for the hypergeometric distribution <0.0001), apoptosis (corrected *P*-value <0.0001) and chromatin organisation (corrected *P*-value <0.0001), while transcription factors of ULK2 seem to be important in homeostatic and immune system-related processes, including glucose homeostasis (corrected *P*-value <0.0001) and response to cytokines (corrected *P*-value <0.0001) (Figure 7).

**Figure 7:**
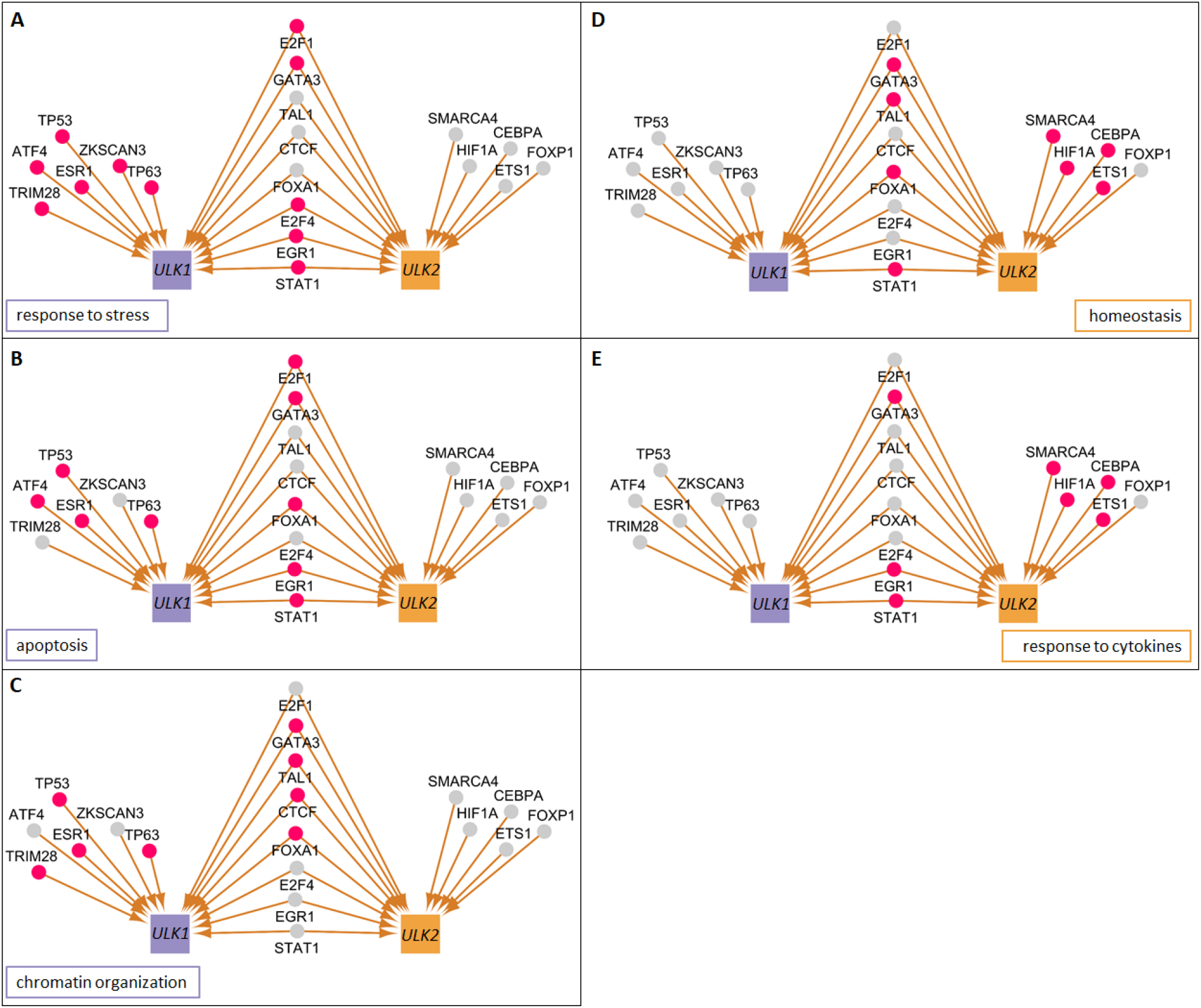
The transcriptional regulators of *ULK1* and *ULK2* and five biological processes with *ULK*-specific regulators. The specific biological processes are labelled on each network, transcription factors involved in the respective biological process are highlighted with pink nodes. A-C) shows processes predominantly specific to regulators of *ULK1*: A) stress-related biological processes; B) apoptosis; C) chromatin organization; while D-E) shows processes predominantly specific to regulators of *ULK2*: D) homeostasis; E) response to cytokines.

## Discussion

Here we report that despite being paralogs and the most similar in their secondary domain structure amongst the ULK homologs, the autophagy induction-related human ULK1 and ULK2 proteins are different in numerous aspects. The appearance of the *ULK1* and *ULK2* genes is the result of the duplication of the yeast *Atg1*. This duplication happened between the Deuterostomes and the Chordates, giving rise to two similar yet functionally different genes. With comprehensive bioinformatics approaches we show that ULK1 and ULK2 have specific protein interactors and transcriptional regulators, involved in distinct biological functions, supporting the specific roles of ULK1 and ULK2. Analysis of the interactors and regulators of ULK1 and ULK2 resulted in the identification of stress response, apoptosis, chromatin organization and catabolic processes as significantly shared among ULK1 interactors, whereas interactors of ULK2 are involved in metabolic pathways, homeostasis and response to cytokines.

As reported in the literature, whole-genome duplication(s) happened throughout vertebrate evolution. The first such event happened at the base of the vertebrates, after the split of *Ciona* from vertebrates 500 million years ago, but prior to the split of fish from tetrapods^18–20^. In concordance with this, we found that the duplication of the *ULK1* ortholog also happened after the split of *Ciona* from vertebrates, so it was likely not a separate event but part of the whole-genome duplication.

As ULK1 and ULK2 both had been identified as part of the autophagy induction complex, the two proteins are often mentioned interchangeably. However, as some studies have already shown, there are functional differences between them, for example they have opposing effects in lipid metabolism^10^. The functional dissimilarity can be a consequence of the different interacting partners and transcriptional regulators that we identified based on databases and manual curation. Regarding the protein-protein interactions, we found 14 common, 11 ULK1-specific and 21 ULK2-specific interactors. We assessed this interactor list with a domain-motif interaction analysis, and identified proteins amongst the experimentally validated ULK1 and ULK2-specific interactors which are in fact predicted to be capable of binding to both ULK1 and ULK2. We note that as we checked the source publication of each interaction we extracted from databases, we found that in many cases only ULK1 or ULK2 was investigated, making the specificity of an interactor to ULK1 or ULK2 questionable. Therefore, we provide a set of proteins that are predicted to bind to both ULK homologs but have not yet been shown *in vitro* to interact for future experimental validation studies. In our analysis we rely on experimentally verified interaction data, hence the uncertainty of the specificity becomes a limitation to our computational study as it could affect the functional gene ontology analysis. Our results are nonetheless supported by the literature, hence we believe our findings can unravel the so far hidden differences of ULK1 and ULK2.

Beside protein-protein interactions we also collected transcriptional regulators of the two *ULK* homologs, and annotated transcription factor binding sites in one or both of the *ULK* homolog promoters. However, as the regulatory connections were mostly determined in high-throughput studies, in most cases we do not know if the specific transcription factors are actually specific to the respective *ULK* gene. One exception is the case of p53. This is because Gao *et al.* analysed the effect of p53 on both *ULK* genes, and found that ectopic expression of p53 results in elevated level of *ULK1*, but not *ULK2^21^*. Interestingly, for p53, we found two transcription factor binding sites on the promoter of *ULK1*, but we also found one binding site along the promoter of *ULK2*. It is possible that because of the two transcription factor binding sites, p53 is more likely to bind to the promoter of *ULK1*, hence affecting its expression. Beside p53, other *ULK1* or *ULK2*-specific transcription factors (ATF4, ESR1, CEBPA, HIF1A, FOXP1) have binding sites on the other *ULK* homolog as well, which makes them a potential pool for further experimental validation.

To investigate the functional differences between ULK1 and ULK2, we analysed their protein interactors and transcriptional regulators for different biological processes and compared the results with the identified ULK1- and ULK2-specific protein motifs. ULK1 and ULK2 had already been found to be partially redundant, as for example double knockdown of *ULK1* and *ULK2* results in a severe reduction of glucose consumption in HCT116 cells, whereas *ULK1* and *ULK2* single knockdowns result in moderate reduction^22^. Nonetheless, *ULK1*, but not *ULK2* knockdown reduces the glucose consumption significantly^22^. This is consistent with our observations that ‘organic substance catabolic processes’ is a significant GO term shared between interactors of ULK1 but not ULK2. In another study, Ro *et. al.* showed that 3T3-L1 cells (a cell line model for adipocytes) do not require *ULK1* for adipogenesis, while knockdown of other autophagy genes, including *ULK2,* suppressed adipogenesis^10^. *ULK2* expression was upregulated in *ULK1* knockdown cells, and *vice versa*, but in *ULK2* knockdown cells upregulation of *ULK1* did not rescue the phenotype. In *ULK2* knockdown cells the expression of *PPARG* and *CEBPA* was decreased too, which also supports our results as CEBPA is one of the *ULK2*-specific transcription factors we highlighted.

We identified that protein interactors of ULK1 but not ULK2 were annotated with the process of stress response, which is in agreement with the results from Wang *et al.*, where they showed that stress granule proteins were enriched in the ULK1 interactome^23^. For creating an ULK1 interactome they used quantitative mass spectrometry and data from the STRING, BioPlex, InWeb, and BioGRID databases. Except for the BioGRID database, these are different databases to the ones used in this study, making their finding an independent evidence contributing to our results. Even though they did not check the interactome of ULK2, they performed experiments with both proteins and showed that ULK1 and ULK2 localized to stress granules. Nonetheless, as interacting partners of ULK1 share the GO term for cellular response to stress, our hypothesis is that ULK1 could have a bigger influence on stress response than ULK2. The results of the functional analysis are further supported by the presence of a phosphorylation site (DOC_CyclinA_RxL_1: Cyclin docking motif), specific to ULK1, on which cyclin/Cdk complexes involved in different biological processes, such as stress response can influence ULK1 (Table 1).

Importantly, our functional analysis suggests that ULK1 and not ULK2 has a specific role in apoptosis and programmed cell death as we found the apoptotic process being one of the significant GO terms that is shared among the transcription factors of ULK1. A crosstalk between apoptosis and autophagy is already well-described, but the difference between ULK1 and ULK2 in the process has not yet been defined^24^. We found ULK1-specific differences in relation to apoptosis on the protein interaction level: we identified an ULK1-specific motif (CLV_C14_Caspase3-7), which is annotated as a site for Caspase-3 and Caspase-7 cleavage. Kim et. al examined another autophagy-related protein, Beclin-1, and found that it is cleaved by caspases^25,26^. Strikingly, Beclin-1 contains the same protein motif we identified on ULK1, confirming that ULK1 could be indeed affected by apoptosis. Moreover, ULK1, but not ULK2, also has a motif (LIG_BIR_III_2) responsible for binding to inhibitor of apoptosis proteins (IAPs). IAPs are known to negatively affect apoptosis^27^, therefore inactivation of these by ULK1 could lead to apoptosis promotion. These two motifs (CLV_C14_Caspase3-7 and LIG_BIR_III_2) could result in a positive feedback for apoptosis, where interaction between ULK1 and IAPs leads to apoptosis, while the presence of caspases promote cleavage of ULK1, hence inhibiting autophagy. Based on our analysis, this feedback is not likely to happen with ULK2. On the contrary, a motif on ULK2 (LIG_EF_ALG2_ABM_2) was previously annotated as being responsible for binding to ALG-2 protein in a Ca2+-dependent manner. ALG-2 has been implicated in ER-stress-induced apoptosis, and shown to be pro-apoptotic^28,29^. This leads us to assume that contrary to ULK1, ULK2 can in fact have a negative effect on apoptosis (Figure 8). Validation of the distinct effects of ULK1 and ULK2 on apoptosis could also shed light on further details of the fine regulation between apoptosis and autophagy.

**Figure 8:**
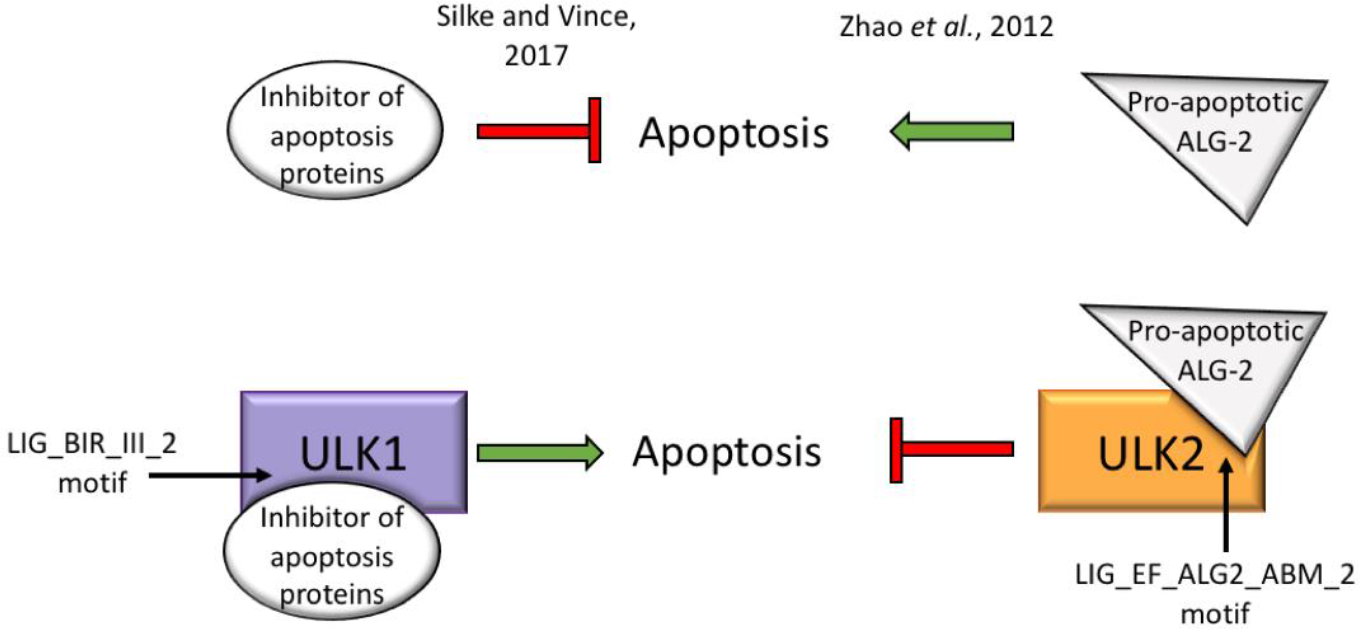
ULK1 and ULK2 could have opposing effects on apoptosis. Based on our analysis ULK1 and ULK2 harbour unique motifs (motifs that are present on only one of them): ULK1 has a LIG_BIR_III_2 motif, which is responsible for binding to inhibitor of apoptosis proteins (IAPs) potentially leading to promotion of apoptosis, while ULK2 has a LIG_EF_ALG2_ABM_2 motif, which can bind the pro-apoptotic ALG-2, resulting in inhibition of apoptosis.

*ULK2*, opposed to *ULK1*, has previously been also shown to be essential for degradation of ubiquitinated proteins and homeostasis in skeletal muscle^16^. These results support our finding that transcription factors of *ULK2*, but not *ULK1,* are annotated to be significant in homeostasis. While our study is not specific for skeletal tissue, we show that *ULK2* definitely has the potential to have a greater impact on tissue homeostasis, which can be especially elevated in tissues with higher expression of *ULK2* than of *ULK1*.

Regarding further specific motifs, ULK2 harbours a TRAF6 binding site (LIG_TRAF6 motif), which is responsible for a response to the tumor necrosis factor receptor (TNFR) superfamily, and direct interaction with various TNF cytokine receptors. The presence of the TRAF6 binding site motif on ULK2 but not on ULK1 underlines the finding that ULK2-specific transcriptional regulators share the function for response to cytokines. This seems relevant in inflammatory diseases, like ulcerative colitis (UC), where the level of cytokines is increased. Interestingly, as we found in two microarray datasets (GSE6731, GSE53306), *ULK2* is downregulated in colon biopsies from inactive UC compared to healthy patients (The lists of differentially expressed genes from the GSE6731 and GSE53306 are included in Supplementary table S1). It is crucial to highlight that while in case of functions shared by ULK1 and ULK2, ULK homologs can have a compensatory effect when one of the homologs is missing, but compensation of specific functions is not possible. In the case of UC, ULK-specific functions can cause an insufficient reaction to cytokines, but also, as ULK1 and ULK2 seem to have opposing effects on apoptosis, the downregulation of *ULK2* can be responsible for the subsequently increased apoptosis, which was previously observed at acute inflammatory sites of patients with inflammatory bowel disease^29^.

In conclusion, with our computational analysis, we have shown that two homologs in the autophagy induction complex, ULK1 and ULK2 likely have specific roles in certain biological processes controlled and mediated by ULK-specific upstream and downstream protein interactors, and transcriptional regulators. In order to understand the consequence of the presented key differences between ULK1- and ULK2-specific regulation of autophagy, experimental studies for disease conditions such as UC should consider analysing these two genes separately.

## Materials and methods

For the comparison of domain structures of the homologs, we used the Pfam website (http://pfam.xfam.org/)^30^. For the annotation of the ULK1 gene duplication event, the dendrogram was adapted from the OMA orthology browser (https://omabrowser.org/)^31^: the branches were collapsed into representative groups and information about the number of homologs in the respective taxa was added. DNA and protein sequences were retrieved from Uniprot (https://www.uniprot.org/)^32^ and their protein alignments were carried out with MUSCLE (https://www.ebi.ac.uk/Tools/msa/muscle/)^33^ with default parameters and ClustalW output format. The protein motifs were downloaded from the Eukaryotic Linear Motif (ELM) resource (http://elm.eu.org/)^34^. The motifs on ULK1 and ULK2 went through a globular domain filtering, structural filtering and context filtering, so we analysed only the accessible motifs that are either outside of globular protein domains or they have an acceptable structural score. The promoter sequences of the human genes ULK1 and ULK2 were retrieved using the “retrieve sequence” function of RSAT (http://www.rsat.eu/)^35^. For the genome assembly of these genes we used the Genome Reference Consortium Human Build 38 patch release 13 (GRCh38.p13)^36,37^. The genome annotation was obtained with the “Full genome build” method (version GENCODE 19), and it contains protein-coding and non-coding genes, splice variants, cDNA, protein sequences and non-coding RNAs. The transcription factor binding site prediction was carried out using the RSAT “matrix-scan” tool and the JASPAR 2018 non-redundant matrices containing the binding profiles of human transcription factors (TFs) (http://jaspar2018.genereg.net/)^38^. Predicted sites with a P-value of <0.0001 were considered to be significant.

Interaction data was downloaded from databases on the 10^th^ of July, 2019. We used the Autophagy Regulatory Network (ARN, http://autophagyregulation.org/)^39^, OmniPath (http://omnipathdb.org/)^40^, TRRUST (version 2, http://www.grnpedia.org/trrust/)^41^, ORegAnno (version 3.0, http://www.oreganno.org/)^42^ and HTRIdb (http://www.lbbc.ibb.unesp.br/htri/)^43^ databases to collect experimentally verified, directed protein-protein (PPI) and transcription factor-target gene interactions for human. Additionally, we used PPI predictions based on the Pfam (http://pfam.xfam.org/)^30^ and ELM (http://elm.eu.org/)^34^ databases. The ARN contains PPI and TF-TG interactions specifically related to autophagy, whereas the other databases are more general, so we filtered the data to obtain a subset of interactions containing ULK1 and ULK2-related connections.

To obtain the significantly shared biological processes of interactors of ULK1 and ULK2, we used the Generic Gene Ontology Term Finder (https://go.princeton.edu/cgi-bin/GOTermFinder)^17^, and we used a cutoff above 30% of the interactors being involved in a certain process. 30% was chosen after examining the distribution of number of interactors and their functions: at 30% of the interactors selected the number of biological processes was optimal. As the GOTermFinder gives significant GO terms as a result, the corrected *P*-value was below 0.01 for all GO terms. Microarray datasets containing biopsy samples from inactive UC and healthy patients (GSE6731, GSE53306) were analysed with the GEO2R function of the Gene Expression Omnibus (https://www.ncbi.nlm.nih.gov/geo/) with default settings. For network visualisation of the ULK1 and ULK2-related interactions we used the Cytoscape program (version 3.7.0, https://cytoscape.org/).

## Supplementary information

**Supplementary Figure S1:**
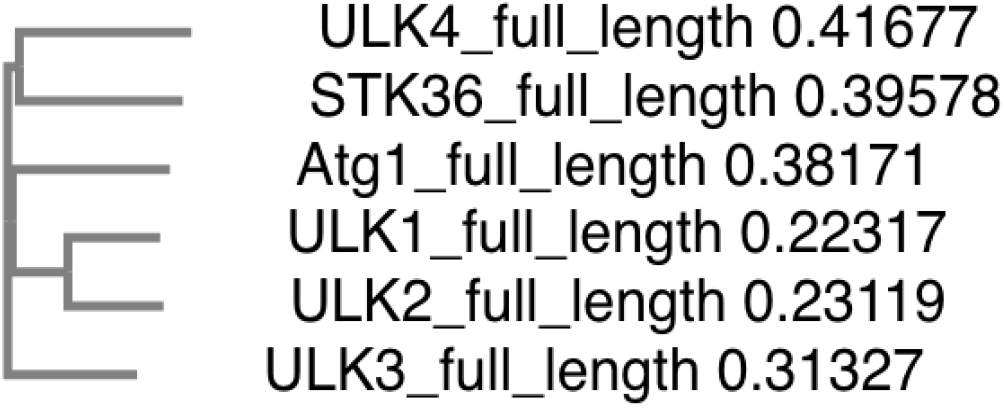
Phylogenetic tree of full the yeast Atg1 protein and its human homologs (complete protein). Based on multiple sequence alignment of the whole length of the proteins, ULK1 and ULK2 are the most similar to each other.

**Supplementary Table S1: Differentially expressed genes from comparing microarray datasets containing biopsy samples from inactive UC and healthy patients (Datasets GSE6731 and GSE53306 from the Gene Expression Omnibus).** Log fold change values were calculated using the built-in GEO2R function of the GEO website. The table contains the calculated median of all values to the same gene (gene symbol), if the adjusted P value was above 0.1. The table contains only genes with abs(med(logFC)) >= 0.585 and the highest adjusted P value was kept.

## Acknowledgement

The authors are grateful for the helpful discussions to the past and present members and visitors of the Haerty and Korcsmaros groups.

## Author contributions

TK and AD worked on the pipeline development; AD, MCRM and MO carried out the protein alignments; AD and MO analysed the phylogenetic tree of ULK1 homologues; AD and MCRM carried out the network analysis and visualisation; LCs worked on the protein-protein interaction prediction; MCRM analysed the protein motifs; PS performed the transcription factor binding site analysis; AD carried out the GO analysis; TK and WH designed and supervised the study. AD, MCRM, WH and TK wrote the manuscript. All authors reviewed the manuscript.

## Data availability

The datasets generated during and/or analysed during the current study are available from the corresponding author.

## Additional information

The authors declare no competing interests.

## References

1. Mizushima, N. & Komatsu, M. Autophagy: renovation of cells and tissues. Cell 147, 728–741 (2011).

2. Lee, E.-J. & Tournier, C. The requirement of uncoordinated 51-like kinase 1 (ULK1) and ULK2 in the regulation of autophagy. Autophagy 7, 689–695 (2011).

3. Alers, S., Löffler, A. S., Wesselborg, S. & Stork, B. Role of AMPK-mTOR-Ulk1/2 in the regulation of autophagy: cross talk, shortcuts, and feedbacks. Mol. Cell. Biol. 32, 2–11 (2012).

4. Kubisch, J. et al. Complex regulation of autophagy in cancer - integrated approaches to discover the networks that hold a double-edged sword. Semin. Cancer Biol. 23, 252–261 (2013).

5. Parzych, K. R. & Klionsky, D. J. An overview of autophagy: morphology, mechanism, and regulation. Antioxid. Redox Signal. 20, 460–473 (2014).

6. Hosokawa, N. et al. Nutrient-dependent mTORC1 association with the ULK1-Atg13-FIP200 complex required for autophagy. Mol. Biol. Cell 20, 1981–1991 (2009).

7. Földvári-Nagy, L., Ari, E., Csermely, P., Korcsmáros, T. & Vellai, T. Starvation-response may not involve Atg1-dependent autophagy induction in non-unikont parasites. Sci. Rep. 4, 5829 (2014).

8. Chan, E. Y. & Tooze, S. A. Evolution of Atg1 function and regulation. Autophagy 5, 758–765 (2009).

9. Lazarus, M. B., Novotny, C. J. & Shokat, K. M. Structure of the human autophagy initiating kinase ULK1 in complex with potent inhibitors. ACS Chem. Biol. 10, 257–261 (2015).

10. Ro, S.-H. et al. Distinct functions of Ulk1 and Ulk2 in the regulation of lipid metabolism in adipocytes. Autophagy 9, 2103–2114 (2013).

11. Shukla, S. et al. Methylation silencing of ULK2, an autophagy gene, is essential for astrocyte transformation and tumor growth. J. Biol. Chem. 289, 22306–22318 (2014).

12. Chan, E. Y. W., Longatti, A., McKnight, N. C. & Tooze, S. A. Kinase-inactivated ULK proteins inhibit autophagy via their conserved C-terminal domains using an Atg13-independent mechanism. Mol. Cell. Biol. 29, 157–171 (2009).

13. Kundu, M. et al. Ulk1 plays a critical role in the autophagic clearance of mitochondria and ribosomes during reticulocyte maturation. Blood 112, 1493–1502 (2008).

14. Cheong, H., Lindsten, T., Wu, J., Lu, C. & Thompson, C. B. Ammonia-induced autophagy is independent of ULK1/ULK2 kinases. Proc Natl Acad Sci USA 108, 11121–11126 (2011).

15. Zachari, M. & Ganley, I. G. The mammalian ULK1 complex and autophagy initiation. Essays Biochem. 61, 585–596 (2017).

16. Fuqua, J. D. et al. ULK2 is essential for degradation of ubiquitinated protein aggregates and homeostasis in skeletal muscle. FASEB J. 33, 11735–11745 (2019).

17. Boyle, E. I. et al. GO::TermFinder--open source software for accessing Gene Ontology information and finding significantly enriched Gene Ontology terms associated with a list of genes. Bioinformatics 20, 3710–3715 (2004).

18. Dehal, P. & Boore, J. L. Two rounds of whole genome duplication in the ancestral vertebrate. PLoS Biol. 3, e314 (2005).

19. Holland, L. Z. & Ocampo Daza, D. A new look at an old question: when did the second whole genome duplication occur in vertebrate evolution? Genome Biol. 19, 209 (2018).

20. Passamaneck, Y. J. & Di Gregorio, A. Ciona intestinalis: chordate development made simple. Dev. Dyn. 233, 1–19 (2005).

21. Gao, W., Shen, Z., Shang, L. & Wang, X. Upregulation of human autophagy-initiation kinase ULK1 by tumor suppressor p53 contributes to DNA-damage-induced cell death. Cell Death Differ. 18, 1598–1607 (2011).

22. Li, T. Y. et al. ULK1/2 constitute a bifurcate node controlling glucose metabolic fluxes in addition to autophagy. Mol. Cell 62, 359–370 (2016).

23. Wang, B. et al. ULK1 and ULK2 regulate stress granule disassembly through phosphorylation and activation of vcp/p97. Mol. Cell 74, 742–757.e8 (2019).

24. Baehrecke, E. H. Autophagy: dual roles in life and death? Nat. Rev. Mol. Cell Biol. 6, 505–510 (2005).

25. Kim, A., Im, M., Yim, N.-H., Kim, T. & Ma, J. Y. A novel herbal medicine, KIOM-C, induces autophagic and apoptotic cell death mediated by activation of JNK and reactive oxygen species in HT1080 human fibrosarcoma cells. PLoS ONE 9, e98703 (2014).

26. Luo, S. & Rubinsztein, D. C. Apoptosis blocks Beclin 1-dependent autophagosome synthesis: an effect rescued by Bcl-xL. Cell Death Differ. 17, 268–277 (2010).

27. Silke, J. & Vince, J. Iaps and cell death. Curr. Top. Microbiol. Immunol. 403, 95–117 (2017).

28. Lacanà, E., Ganjei, J. K., Vito, P. & D’Adamio, L. Dissociation of apoptosis and activation of IL-1beta-converting enzyme/Ced-3 proteases by ALG-2 and the truncated Alzheimer’s gene ALG-3. J. Immunol. 158, 5129–5135 (1997).

29. Zhao, Q., Yu, Y., Zuo, X., Dong, Y. & Li, Y. Milk fat globule-epidermal growth factor 8 is decreased in intestinal epithelium of ulcerative colitis patients and thereby causes increased apoptosis and impaired wound healing. Mol. Med. 18, 497–506 (2012).

30. El-Gebali, S. et al. The Pfam protein families database in 2019. Nucleic Acids Res. 47, D427–D432 (2019).

31. Altenhoff, A. M. et al. The OMA orthology database in 2018: retrieving evolutionary relationships among all domains of life through richer web and programmatic interfaces. Nucleic Acids Res. 46, D477–D485 (2018).

32. UniProt Consortium, T. UniProt: the universal protein knowledgebase. Nucleic Acids Res. 46, 2699 (2018).

33. Madeira, F. et al. The EMBL-EBI search and sequence analysis tools APIs in 2019. Nucleic Acids Res. 47, W636–W641 (2019).

34. Gouw, M. et al. The eukaryotic linear motif resource - 2018 update. Nucleic Acids Res. 46, D428–D434 (2018).

35. Nguyen, N. T. T. et al. RSAT 2018: regulatory sequence analysis tools 20th anniversary. Nucleic Acids Res. 46, W209–W214 (2018).

36. Church, D. M. et al. Modernizing reference genome assemblies. PLoS Biol. 9, e1001091 (2011).

37. Schneider, V. A. et al. Evaluation of GRCh38 and de novo haploid genome assemblies demonstrates the enduring quality of the reference assembly. Genome Res. 27, 849–864 (2017).

38. Khan, A. et al. JASPAR 2018: update of the open-access database of transcription factor binding profiles and its web framework. Nucleic Acids Res. 46, D260–D266 (2018).

39. Türei, D. et al. Autophagy Regulatory Network - a systems-level bioinformatics resource for studying the mechanism and regulation of autophagy. Autophagy 11, 155–165 (2015).

40. Türei, D., Korcsmáros, T. & Saez-Rodriguez, J. OmniPath: guidelines and gateway for literature-curated signaling pathway resources. Nat. Methods 13, 966–967 (2016).

41. Han, H. et al. TRRUST v2: an expanded reference database of human and mouse transcriptional regulatory interactions. Nucleic Acids Res. 46, D380–D386 (2018).

42. Lesurf, R. et al. ORegAnno 3.0: a community-driven resource for curated regulatory annotation. Nucleic Acids Res. 44, D126–32 (2016).

43. Bovolenta, L. A., Acencio, M. L. & Lemke, N. HTRIdb: an open-access database for experimentally verified human transcriptional regulation interactions. BMC Genomics 13, 405 (2012).

